# A mechanism-aware stoichiometry platform resolves functional viral thresholds and induces antiviral hypersensitivity

**DOI:** 10.64898/2026.02.09.704886

**Authors:** Gabe Clinger, Haley Durden, Aaron Daurelle, Benjamin Preece, Wiley Peppel, Rodrigo Gallegos, Abdul A. Waheed, Nicole Bohne, Brian MacArthur, Abby Peterson, Ahmet Yildiz, Eric O. Freed, Saveez Saffarian

**Author notes:** Authors contributed equally (Gabe Clinger, Haley Durden, Aaron Daurelle).

## Abstract

Antiviral discovery is frequently hindered by a ‘stoichiometric blind spot’—a lack of resolution regarding the functional density of viral enzymes required for replication. Traditional screening pipelines rely on target-based or phenotypic assays that cannot distinguish between simple molecular binding and the crossing of a functional ‘stoichiometric cliff.’ Here, we present a mechanism-aware platform that resolves these enzymatic requirements with single-virion precision. By integrating quantitative cryo-electron microscopy and genomic validation with Monte Carlo modeling, we map the stoichiometric landscapes of HIV-1 Protease (PR) and Reverse Transcriptase (RT). We uncover a striking disparity in enzymatic demand: a high-redundancy ‘buffering capacity’ for PR (∼40 monomers) contrasted with a high-threshold requirement for RT (∼95 subunits). We demonstrate that by systematically ‘de-buffering’ the virion, our platform induces a state of antiviral hypersensitivity, enabling the detection of therapeutic activity in novel and clinical inhibitors that remain invisible to traditional workflows. Furthermore, this multiplexed profiling enables de novo target identification, as inhibitors trigger failure exclusively on their respective stoichiometric arms. This platform provides a deterministic roadmap for de-risking drug discovery and identifying viral sub-stoichiometric vulnerabilities.

## Introduction

The replication of Human Immunodeficiency Virus type 1 (HIV-1) is a tightly orchestrated process governed by a limited set of viral enzymes packaged within the virion^1^. Central to this process are the products of the *pol* gene: Protease (PR), Reverse Transcriptase (RT), and Integrase (IN). Each plays a distinct, non-redundant role in the viral replication cycle^2^. PR is responsible for the proteolytic processing of Gag and Gag-Pol polyproteins, a process known as maturation that transforms an immature, non-infectious particle into a mature virion containing a conical core^3^. Once a new cell is entered, RT converts the viral single-stranded RNA genome into double-stranded DNA, which is subsequently inserted into the host genome by IN^4^. In a remarkable feat of biological engineering, HIV-1 ensures the equimolar incorporation of these enzymes by packaging approximately 100 copies of the Gag-Pol polyprotein into every virion^5^. A ∼20:1 ratio of Gag:Gag-Pol expression in the infected cell is ensured by a ribosomal frameshifting event during Gag translation that results in the synthesis of Gag-Pol.

Despite the packaging of stoichiometric amounts of PR, RT, and IN during virus assembly, it has remained unclear whether the virus actually *requires* every packaged enzyme to successfully replicate^6^. In the pharmaceutical industry, the search for inhibitors of these enzymes has traditionally relied on two pillars: target-based high-throughput screening (HTS) and phenotypic drug discovery (PDD)^7^. Target-based HTS identifies high-affinity binders using purified proteins in isolation, often ignoring the crowded, redundant environment of the native virion^8,9^. Conversely, phenotypic screening measures total viral inhibition in live cells but operates as a “black box,” unable to resolve the specific “stoichiometric cliff”: the minimum number of functional enzymes required for replication^10^.

The discovery of Lenacapavir (GS-6207), a first-in-class capsid inhibitor, highlights this technological gap. Although it achieved unprecedented picomolar potency by interfering with both nuclear entry and virion maturation, its multistage mechanism was only fully understood through years of post-hoc structural analysis^11,12^. This retrospective discovery process underscores a fundamental limitation in current drug development: the industry lacks a ‘mechanism-aware’ platform capable of resolving functional thresholds in real-time during lead optimization. Because standard phenotypic assays operate as a ‘black box’ regarding the molecular count required for inhibition^10^, they remain effectively ‘stoichiometrically blind.’ Consequently, the natural enzymatic redundancy of the virion can mask a drug’s efficacy until a critical threshold is crossed—a phenomenon that hides potential leads and complicates the optimization of first-in-class inhibitors.

Here, we present a quantitative stoichiometry platform that resolves these functional requirements with single-virion precision. By combining defined stoichiometric perturbations with Monte Carlo modeling, we quantify the minimal requirements for HIV-1 enzymes. Our results show a striking disparity in the number of enzyme molecules required for productive infection: while maturation proceeds normally with only ∼40 of the ∼100 packaged PR monomers, reverse transcription requires a nearly full complement of ∼95 functional RT subunits.

This framework uncovers a significant “buffering capacity” in HIV-1 maturation that mutes the observable effects of low-concentration inhibitors in standard assays. By systematically removing this buffer, we demonstrate a state of antiviral hypersensitivity that allows for the detection of therapeutic activity such as that of the novel inhibitor PDDC^13^ at concentrations that would remain invisible to traditional pharmaceutical workflows. This platform provides a generalizable roadmap for de-risking drug discovery by mapping the hidden vulnerabilities of viral replication.

## Results

### A Multi-Modal Platform for Stoichiometric Resolution of the HIV-1 Payload

To establish a “mechanism-aware” drug discovery tool, we first developed a platform capable of both manipulating and quantifying the enzymatic architecture of individual HIV-1 virions. We utilized a co-assembly strategy, transfecting cells with varying ratios of wild-type (WT) and catalytically inactive mutant (mut) Gag-Pol plasmids to produce virion populations with precise enzymatic gradients (Fig. 1a-c).

**Figure 1.**
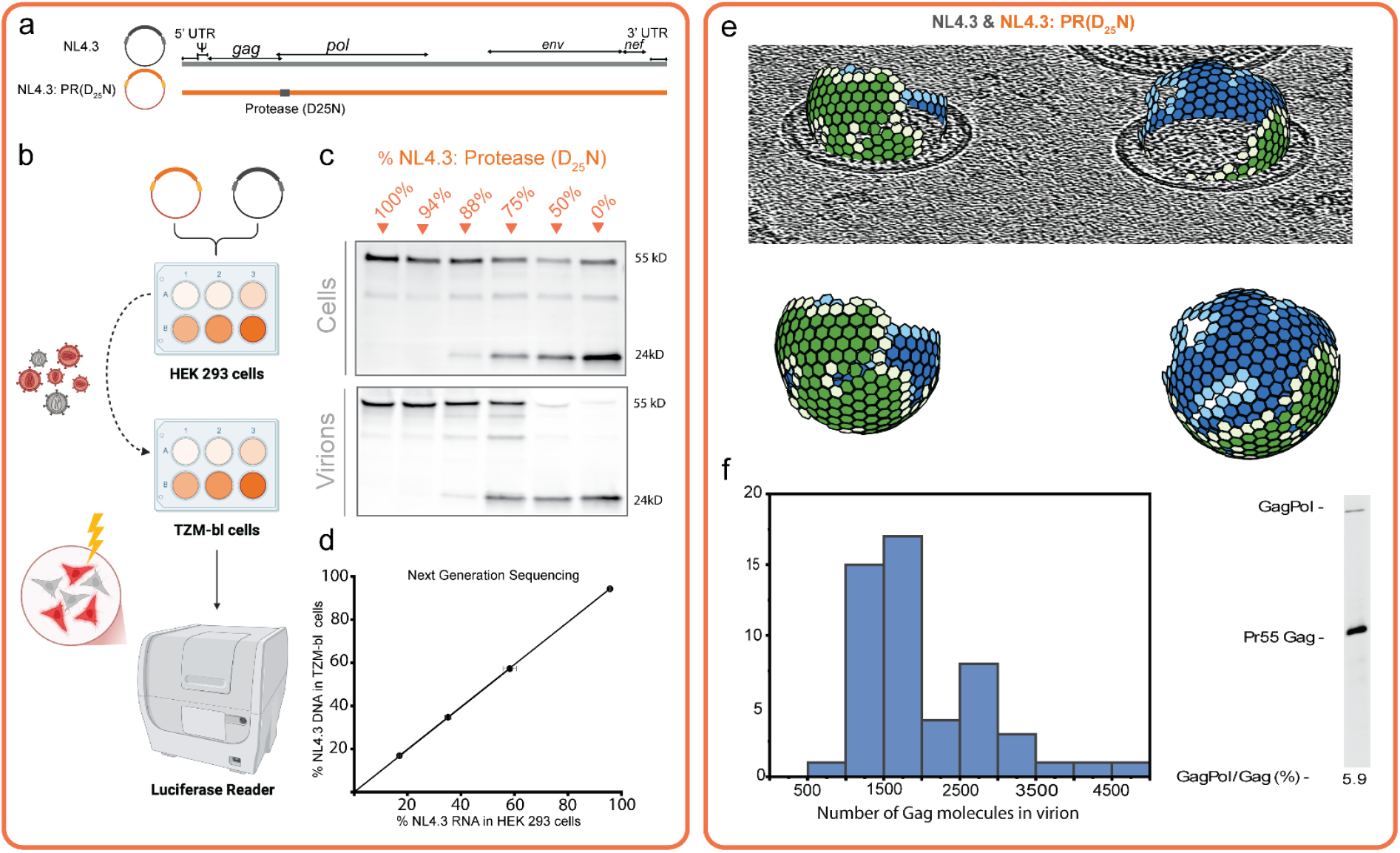
A quantitative platform for titrating viral enzyme stoichiometry in single HIV-1 virions. (a) Architecture of the co-assembly system. Schematic of the wild-type HIV-1 molecular clone (NL4.3, black) and the enzymatically inactive mutant clone (NL4.3: PR(D_25_N), orange). The mutant carries a single-point mutation in the PR active site, rendering it catalytically null while maintaining structural assembly competence. (b) Workflow for stoichiometric titration and infectivity profiling. HEK 293T producer cells are co-transfected with varying ratios of WT and mutant plasmids to produce “hybrid” virions with controlled enzymatic payloads. Resulting virions are harvested to infect TZM-bl reporter cells, where luciferase activity and next-generation sequencing (NGS) provide a readout of infectivity. (c) PR processing across a stoichiometric gradient. Immunoblots of cell lysates (top) and purified virions (bottom) showing the processing of Gag (p55) to mature CA (p24) as a function of the percentage of WT PR (d) Preservation of genetic ratios during assembly. NGS analysis confirms a 1:1 correlation between the percentage of NL4.3 RNA in producer cells and the resulting DNA in target cells, validating that the plasmid ratio accurately predicts the intravirion RNA stoichiometry. (e) Single-virion structural analysis via cryo-ET. Top: Representative cryo-electron tomogram slice of hybrid HIV-1 virions. Bottom: 3D subtomogram averages showing the lattice structure of the Gag shell, confirming that stoichiometric titration does not disrupt the fundamental lattice assembly. (f) Quantifying virion composition. Left: Histogram derived from Cryo-ET and mass spectrometry showing the distribution of Gag molecules per virion (mean ∼2500 copies). Right: Representative western blot and densitometry quantifying the GagPol-to-Gag ratio at 5.9%, establishing the baseline enzymatic payload for the platform.

The utility of this platform rests on two rigorous validation pillars. First, we performed quantitative Cryo-Electron Microscopy (Cryo-EM) to establish the absolute physical payload of the virus. By resolving the density of the viral lattice, we confirmed that HIV-1 packages a remarkably consistent average of ∼100 Gag-Pol molecules per virion (Fig. 1e-f). This physical count provides the essential “N” value for our subsequent mathematical modeling. Second, we used Next-Generation Sequencing (NGS) to verify that the genomic RNA ratios within the harvested virions linearly matched the input plasmid ratios across the entire titration range (Fig. 1d).

Together, these data demonstrate that the platform allows us to treat the virion as a “programmable” biological reactor. With the average number of Gag-Pol molecules per virion fixed at ∼100 molecules, we can now precisely map the “stoichiometric cliff”—the exact point where the number of molecules of an enzyme falls below the threshold required for successful infection.

### HIV-1 Maturation Proceeds with Substantial PR Redundancy

Having established that each HIV-1 virion contains an average of 100 Gag-Pol molecules (Fig. 1), we utilized our platform to determine the minimal number of functional PR monomers required for particle infectivity. HIV-1 PR is responsible for the systematic cleavage of the Gag and Gag-Pol polyproteins, a prerequisite for the formation of the infectious conical core. To probe the stoichiometric threshold of this process, we generated virion populations by co-transfecting WT Gag-Pol with a catalytically inactive D25N mutant across a wide range of ratios.

We observed that viral infectivity remained remarkably stable even as the fraction of active PR was significantly reduced. The resulting infectivity curve followed a shallow decay, maintaining >80% of wild-type infectivity even when the active PR fraction was halved (Fig. 2A). This resilience suggests that HIV-1 maturation possesses a significant “buffering capacity,” where the virus packages an excess of PR to ensure maturation success under sub-optimal conditions. An alternative interpretation is that 100 molecules of Gag-Pol are packaged because the virus needs 100 molecules of RT or IN. The excess PR is thus just a bystander effect; devising a scenario to package less PR than RT or IN is not worth the trouble.

**Figure 2.**
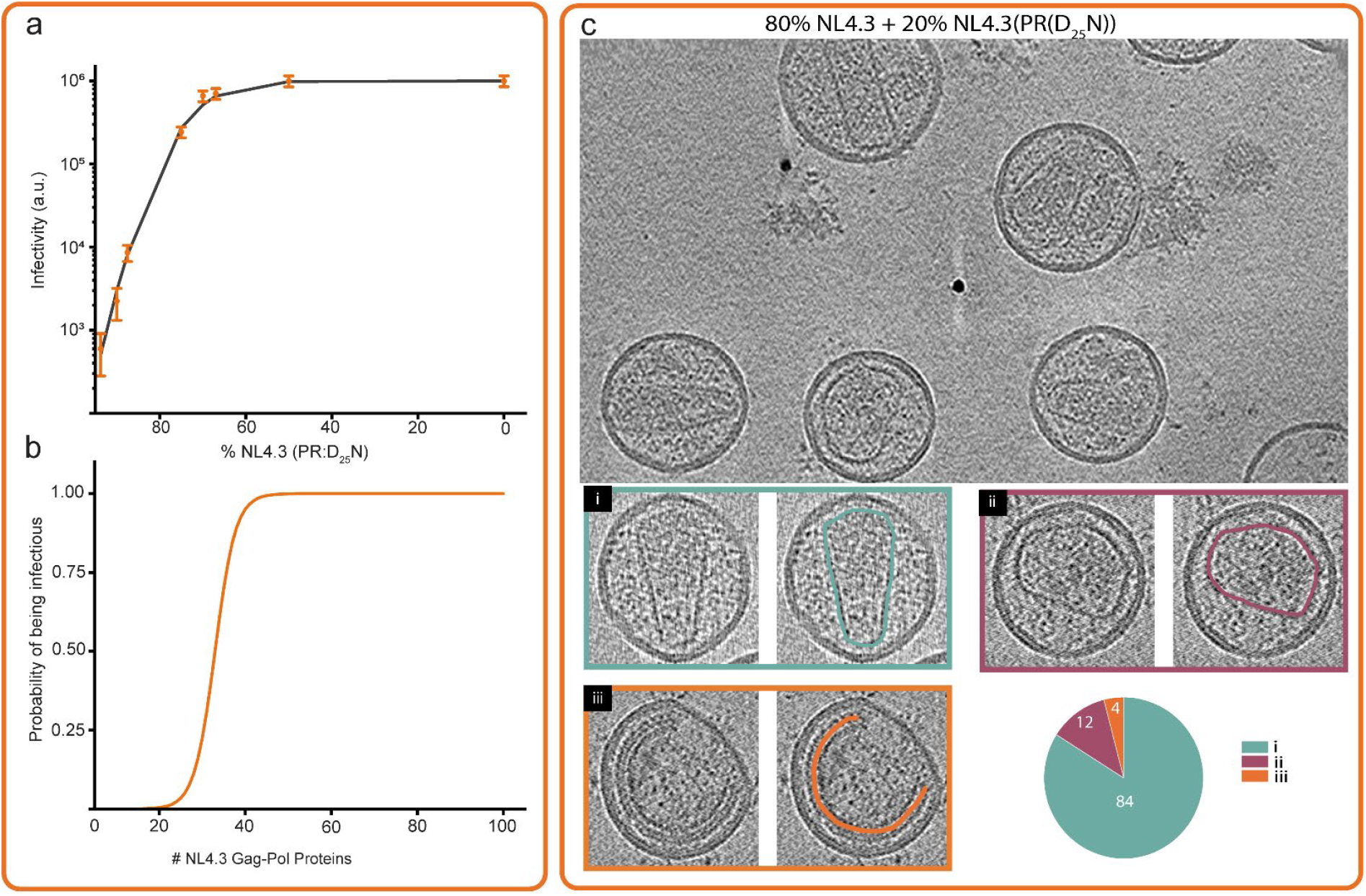
Determining the stoichiometric threshold for HIV-1 infectivity. (a) Titration reveals a nonlinear relationship between inactive PR subunits and particle infectivity. Viral infectivity (measured in arbitrary luciferase units) plotted as a function of the percentage of PR-defective mutant (NL4.3: PR(D_25_N)) incorporated into the virion. The plateau from 0% to 50% mutant indicates a high degree of enzymatic redundancy (“buffering capacity”), followed by a sharp “stoichiometric cliff” where infectivity drops by three orders of magnitude as mutant levels exceed 75%. Error bars represent standard deviation from three independent replicates. (b) Mathematical derivation of the minimal functional payload. A Monte Carlo simulation model, constrained by the experimental data in (a) and the physical virion parameters in Fig. 1f, calculates the probability of a single virion being infectious as a function of the absolute number of wild-type Gag-Pol proteins. The sigmoidal transition identifies a critical threshold of ∼40 Gag-Pol molecules (PR monomers) required for the virus to be infectious. (c) Morphological validation of the maturation threshold via Cryo-ET. Top: Representative cryo-electron tomogram of a hybrid virion population (80% NL4.3 + 20% NL4.3: PR(D_25_N)). Bottom: Categorization of maturation states based on core morphology: (i) mature virions with characteristic conical capsids (green), (ii) defective core architectures eccentric (magenta) (iii) immature-like particles (orange). The pie chart quantifies the distribution, Interestingly, at 80% ratio, the infectivity is not affected while a significant percentage of particles are defective.

To transform these phenotypic observations into discrete molecular counts, we applied our Monte Carlo simulation to the experimental data, assuming that PR functions as an obligate dimer. The model calculates the probability that a virion containing N total Gag-Pol molecules will incorporate at least k functional subunits. The best fit to the experimental PR titration curve yielded a requirement of 40 active monomers (Fig. 2b). This indicates that only ∼40% of the packaged PR payload is strictly necessary to trigger the structural transitions required for maturation.

This finding demonstrates that even with PR where traditionally its observed that replication is severely impacted by inhibitors at low concentration, the viral particles package more PR than what they need. By defining this “stoichiometric cliff” at 40 monomers, our platform identifies the specific degree of inhibition a therapeutic must achieve to effectively block particle infectivity.

### Reverse Transcription is Stoichiometrically Hypersensitive to Active-Site Loss

While the virus possesses significant functional redundancy for maturation, we hypothesized that subsequent replication steps might impose stricter stoichiometric requirements. To test this, we applied our platform to quantify the functional threshold for RT, the enzyme responsible for converting the viral RNA genome into double-stranded DNA. We generated virions by co-transfecting WT Gag-Pol with an RT active-site mutant (D_185_N/D_186_N) across the same titration range used for PR.

In sharp contrast to the shallow decay observed for PR, the loss of active RT subunits resulted in a steep, non-linear collapse of viral infectivity (Fig. 3A). Even minor reductions in the fraction of functional RT led to a profound loss of replication fitness, suggesting that the “buffering capacity” observed during maturation is entirely absent during the reverse transcription phase.

**Figure 3.**
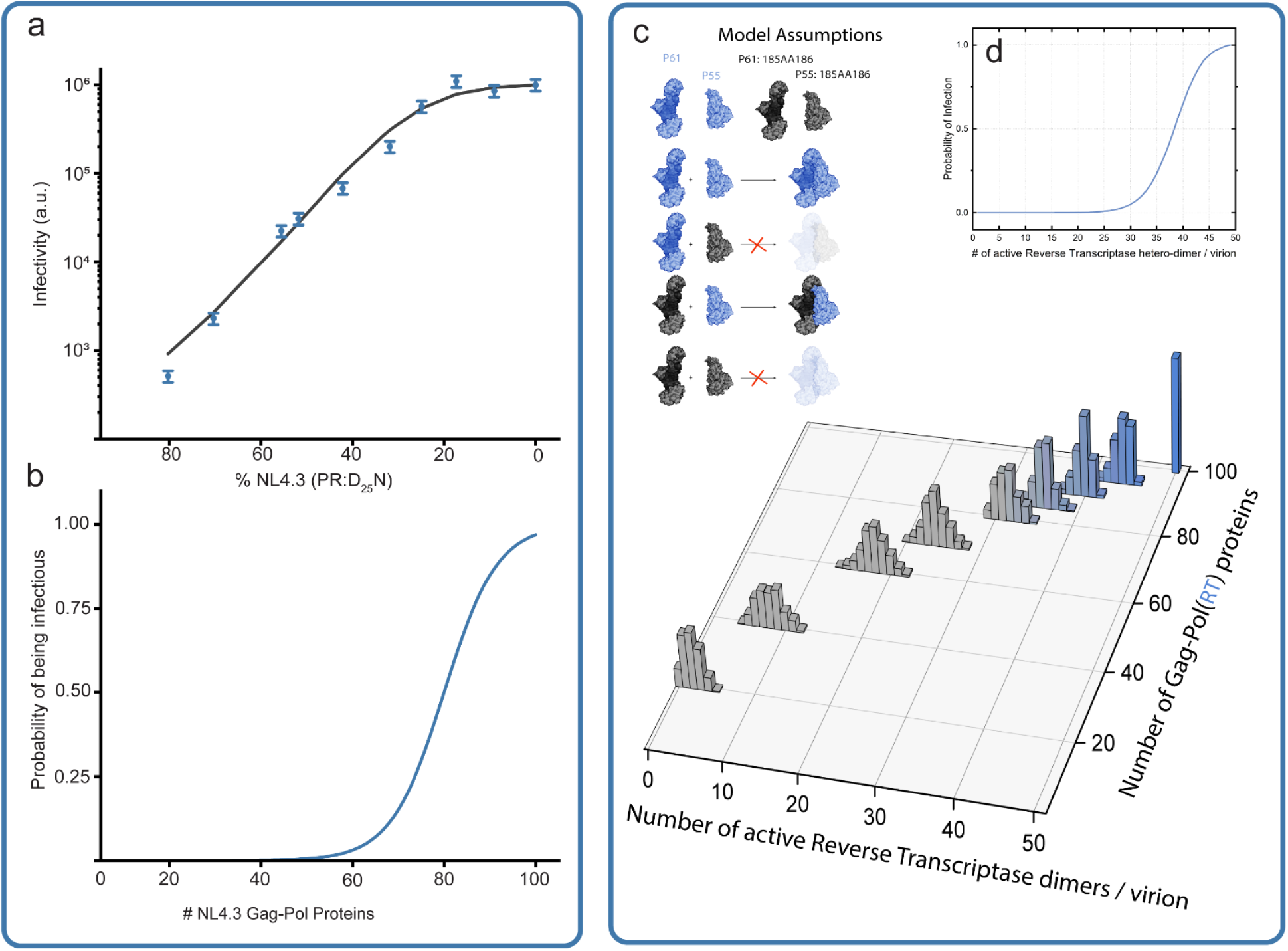
High stoichiometric demand and assembly requirements for HIV-1 RT. (a) Infectivity titration of RT activity. Viral infectivity (arbitrary luciferase units) is plotted against the percentage of wild-type NL4.3 (incorporating active RT) relative to an RT-defective mutant (NL4.3: PR(D_25_N) backbone with RT-inactivating mutations). In contrast to the PR titration (Fig. 2a), RT activity shows a more linear and immediate decline as WT Gag-Pol levels are reduced, indicating a significantly lower “buffering capacity” for reverse transcription. (b) Stoichiometric model of RT functional requirements. Monte Carlo simulation results showing the probability of a single virion being infectious as a function of the absolute number of wild-type Gag-Pol molecules packaged. The sigmoidal transition identifies a high functional threshold, with ∼95 wild-type Gag-Pol molecules required for a 50% probability of infection—nearly the entire natural payload of the virion. (c) Modeling the assembly of functional RT heterodimers. Top left: Model assumptions for the stochastic assembly of p66 and p51 subunits to form active RT heterodimers within the maturing virion core. Bottom: 3D histogram showing the predicted distribution of active RT heterodimers per virion across a range of total packaged Gag-Pol (RT) proteins. The model accounts for subunit availability and the probability of successful dimerization during maturation. (d) Probability of infection relative to active RT heterodimers. Derived functional curve showing the probability of infection based on the absolute number of active RT heterodimers per virion. The model predicts that reverse transcription is a highly inefficient process at the single-virion level, requiring approximately 40–50 active heterodimers to ensure successful replication, representing a critical “stoichiometric bottleneck.”

To determine the exact molecular requirement, we fitted the titration data to our Monte Carlo model. The best-fit simulation revealed a requirement of 95 functional subunits (Fig. 3B). Given our earlier physical quantification of 100 packaged Gag-Pol molecules (Fig. 1), this result indicates that nearly the entire packaged RT payload must be catalytically active to successfully complete the synthesis of the viral cDNA.

This “high-threshold” requirement defines RT as a point of extreme stoichiometric vulnerability. While the virus can tolerate a 60% loss of PR activity, it cannot survive even a 10% loss of RT activity. This finding provides a mechanistic explanation for why RT remains one of the most effective targets for clinical intervention: unlike PR, the virus has no enzymatic “safety net” to buffer against RT inhibitors. In addition, modern PIs are extremely potent but lead to adverse side effects in many patients. Our platform thus identifies RT as a high-value target for “sub-stoichiometric” inhibition, where blocking just a few molecules per virion is sufficient to trigger complete replication failure.

### De Novo Target Identification and Potency Resolution via Multiplexed Stoichiometric Profiling

A central challenge in antiviral discovery is the “target deconvolution” of hits identified through phenotypic screening. To demonstrate that our platform provides simultaneous potency resolution and target identification (Target ID), we performed a multiplexed stoichiometric profile of three chemically distinct protease inhibitors: the clinical standards Darunavir and Saquinavir, and the novel compound PDDC which Inhibits neutral sphingomyelinase 2 and blocks virions maturation. Each compound was applied to both the PR-titrated and RT-titrated virion populations.

The platform provided an unambiguous de novo identification of the inhibitors’ targets. Across all three compounds, we observed a profound “stoichiometric cliff” exclusively on the PR-titration arm, where reducing the functional payload to the 40-threshold resulted in a dramatic leftward shift in infectivity (Fig. 4A). In stark contrast, the RT-titration arm remained entirely unaffected, with dose-response curves for all three inhibitors remaining within background signal levels, regardless of RT occupancy (Fig. 4B). This internal negative control demonstrates that the platform can distinguish between off-target effects and mechanism-specific inhibition with high fidelity.

**Figure 4.**
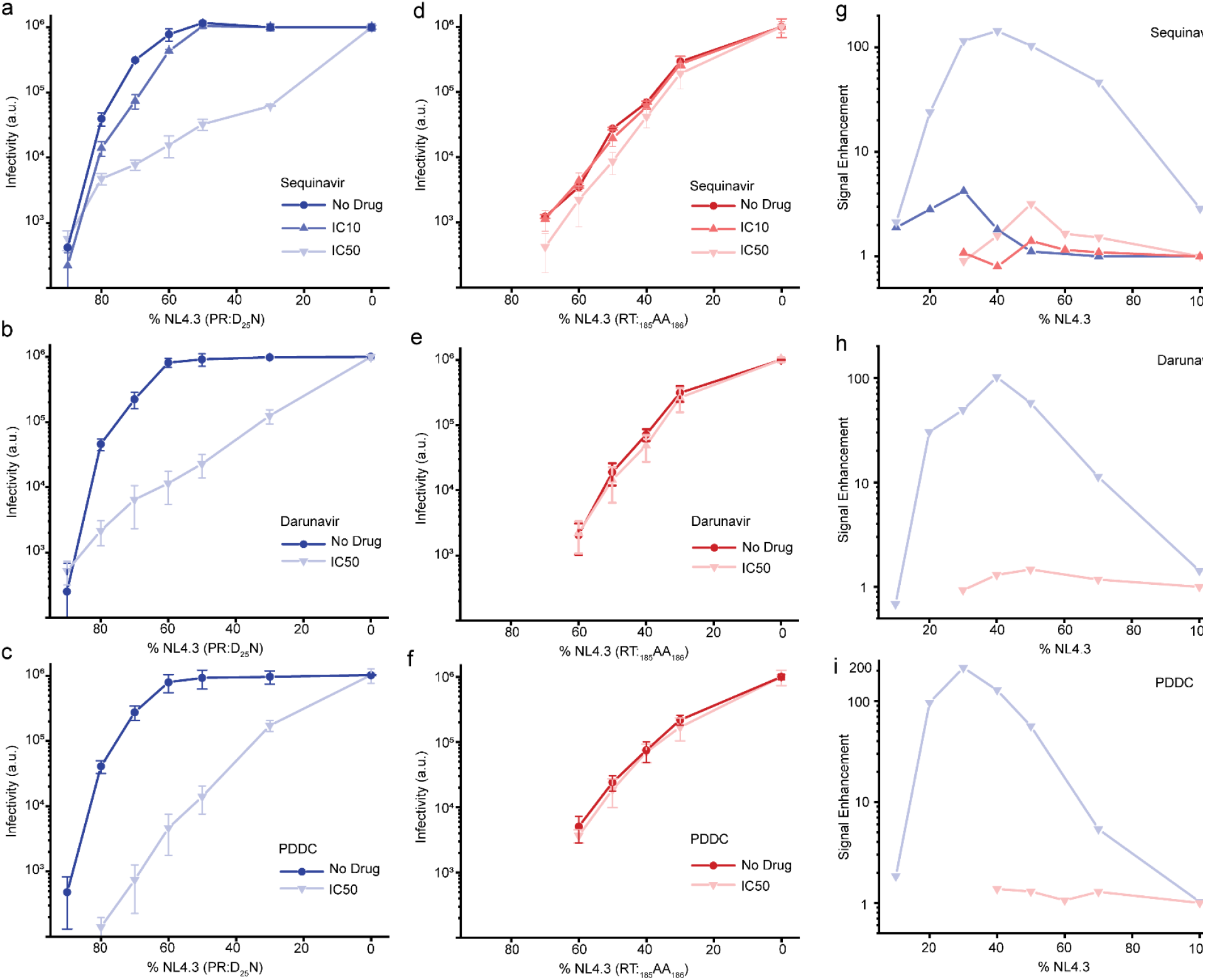
Stoichiometric sensitization reveals antiviral hypersensitivity and drug mechanism of action. a–c, Protease inhibitor (PI) sensitivity under stoichiometric stress. Infectivity profiles (luciferase a.u.) of hybrid virions titrated with the PR-defective mutant NL4.3(PR:D_25_N) in the presence of Saquinavir (a), Darunavir (b), and the experimental compound PDDC (c). While wild-type virions (0% mutant) show minimal response to drugs at IC_10_ or IC_50_ concentrations due to high enzymatic redundancy, reducing the functional PR payload (increasing % mutant) induces a state of hypersensitivity, leading to a precipitous loss of infectivity. d–f, Reverse Transcriptase (RT) inhibitor profiles under stoichiometric stress. Parallel titration experiments using the RT-defective mutant NL4.3(RT_185_AA_186_) in the presence of the same compounds (Saquinavir, Darunavir, PDDC). In sharp contrast to the PR profiles, stoichiometric reduction of RT does not significantly enhance the potency of these PIs, confirming that the “hypersensitivity effect” is target-specific and limited to the enzymatic pathway being inhibited. g–i, Quantification of signal enhancement. Signal enhancement (defined as the ratio of drug-induced inhibition in sensitized vs. wild-type virions) plotted as a function of the wild-type plasmid percentage for Saquinavir (g), Darunavir (h), and PDDC (i). Peaks in signal enhancement (up to ∼200-fold for PDDC) identify the optimal “diagnostic window” where stoichiometric titration reveals therapeutic efficacy that is undetectable in standard phenotypic screens. Blue curves indicate protease-sensitized virions; red curves indicate RT-sensitized virions.

Furthermore, the “signal-to-background” resolution of the PR-arm was strikingly large for all three inhibitors. By “de-buffering” the virions—removing the redundant PR molecules that normally mask drug activity—we amplified the inhibitory signal to a level that revealed sub-stoichiometric potency (Fig. 4C). This hypersensitivity allows the platform to function like a phenotypic screen that “self-deconvolves,” providing immediate Target ID while simultaneously de-masking low-concentration activity that would be lost to the “black box” of traditional bulk assays.

These results suggest that our platform could be utilized to screen uncharacterized chemical libraries, providing a “mechanism-first” discovery engine where the target and the stoichiometric vulnerability are identified in a single step.

## Discussion

We have demonstrated a mechanism-aware platform that resolves the functional stoichiometry of viral enzymes with single-virion precision. By integrating quantitative Cryo-EM, genomic validation, and Monte Carlo modeling, we identified a profound disparity in the enzymatic requirements of HIV-1: a high-redundancy “buffering capacity” for Protease (∼40 monomers) contrasted against a high-threshold hypersensitivity for RT (∼95 subunits). We acknowledge that the steep stoichiometric threshold observed for RT may partially stem from a dominant-negative effect, distinct from simple vacancy. Unlike PR, which dimerizes transiently, RT functions as a stable p66/p51 heterodimer. It is plausible that heterodimers containing mutant subunits retain template-binding affinity despite catalytic inertness, thereby occluding the viral genome from functional enzymes^14^. However, in the context of therapeutic discovery, this distinction reinforces the vulnerability of the system: whether driven by lack of redundancy or competitive occlusion, the RT complex displays an extreme structural intolerance to ‘poisoning’ by non-functional units. This confirms that sub-stoichiometric inhibitors—which mimic this dominant-negative state—face a much lower barrier to efficacy than those targeting high-redundancy enzymes like PR. This stoichiometric mapping provides additional quantitative explanation for why certain drug classes face higher biological hurdles than others, which helps in designing a better roadmap for identifying “stoichiometric cliffs” across diverse viral architectures.

The identification of enzymatic buffering has significant implications for the pharmaceutical industry’s reliance on “black-box” phenotypic and target-based HTS. As demonstrated by the discovery of Lenacapavir, traditional pipelines often lack the resolution to identify the exact target and stoichiometry required for inhibition during early lead optimization. Our platform addresses this by enabling a state of antiviral hypersensitivity. By “de-buffering” the virion, we reveal potent inhibitory signals for compounds like PDDC, Darunavir, and Saquinavir that are otherwise masked by the virus’s natural safety net. This hypersensitivity effectively lowers the detection limit for new chemotypes, allowing for the “rescue” of potent leads that would be discarded in standard bulk assays.

Perhaps the most disruptive feature of the platform is its capacity for de novo Target Identification (Target ID). As shown in our multiplexed profiling (Fig. 4), the platform acts as a self-deconvolving screen: inhibitors trigger a stoichiometric cliff only on their specific target arm (PR), while the non-target arm (RT) remains at background levels. This eliminates the “deconvolution bottleneck” that typically follows phenotypic discovery, providing simultaneous data on potency, mechanism, and target identity in a single assay.

While validated here using HIV-1, this platform may be inherently generalizable. The stoichiometric principles revealed—redundancy versus high-threshold requirements—are likely conserved across other enveloped viruses, including Influenza A and VSV. In the context of pandemic preparedness, the ability to rapidly map the stoichiometric vulnerabilities of an emerging pathogen would allow for the immediate “de-risking” of existing drug libraries and the accelerated design of sub-stoichiometric inhibitors.

In conclusion, our results transition antiviral discovery from a stochastic search for binders to a deterministic mapping of functional failure points. By resolving the hidden stoichiometry of the virion, the Revirion platform provides the high-resolution diagnostic necessary to meet the increasing demand for precision antivirals in both human and animal health.

## Materials and methods

To investigate the co-assembly of HIV-1 virions and its impact on infectivity, we employed a well-established tissue culture system. Specifically, HEK293 cells were utilized for robust production of viral particles following transfection with the pNL4.3 proviral DNA backbone. Viral infectivity was quantitatively assessed using TZM-bl cells, a HeLa cell derivative engineered to express CD4, CCR5, and a Tat-responsive firefly luciferase reporter gene, enabling a direct measurement of infection efficiency. The infectivity observed in this study is therefore defined as the ability of the generated virions to infect TZM-bl cells and induce Tat-dependent luciferase expression.

Our primary objective was to determine how the ratio of wild-type to mutant Gag-Pol polyprotein expression, arising from the co-transfection of pNL4.3 and the respective mutant backbones, affects the overall infectivity of the assembled virions. While a simplified view might suggest a direct correlation between the proportion of wild-type components and infectivity, the assembly process in transfected HEK293 cells is inherently complex. Each cell within the transfected population takes up a variable number of plasmid molecules, leading to a heterogeneous expression of wild-type and mutant viral proteins within individual producer cells. Consequently, the observed infectivity of the viral population is a composite outcome of virions assembled with varying ratios of functional and non-functional components. Accurate modeling of the infectivity data therefore necessitates a quantitative understanding of the distribution of pNL4.3 plasmid uptake and subsequent protein expression across the HEK293 cell population. We next describe the method employed to measure this distribution.

### Co-assembly Experiments: Cell maintenance and transfections for infectivity experiments

Human embryonic kidney (HEK) 293 cell lines were maintained in T-25 flasks using TrypLE Express Enzyme (Gibco) and DMEM with L-Glutamine, 4.5g/L glucose and sodium pyruvate (Corning) supplemented with 10% fetal bovine serum (Gibco). Cells were incubated at 37°C. Once over 90% confluent, the HEK 293 cells were plated onto 6-well cell culture plates with 2 ml of medium per well. Experiments were performed in sets of 6, with each well containing a different ratio of HIV-1 gRNA (pNL4.3) plasmid DNA and a plasmid containing a single-point mutation. Each well was transfected with 8ng total of plasmid DNA. These transfections were performed using 100ul Opti-MEM Reduced Serum Medium (Gibco) and 8ul of Lipofectamine 2000 (Thermo Fisher Scientific) for each plate. The cells were incubated at 37°C for 36-48 hrs.

### Co-assembly Experiments: Harvesting virions and cell extracts for infectivity and western blot assays

To harvest virions for western blot analysis and luciferase assays, cells and supernatants were collected from each well on our 6-well plate. The contents of the plate plate were centrifuged at 5,000 RPM (Ample Scientific Champion S-50D centrifuge) for 10 min to separate the cells from the supernatant, and then the supernatant was filtered using a 0.22um syringe filter (Celltreat). Before proceeding with further purification of the virions, 1ml of this filtered supernatant was set aside to be used in a luciferase assay. The cell pellets were washed with PBS (Quality Biological) and then resuspended in 100ul of RIPA lysis buffer (Santa Cruz Biotechnology) containing PR inhibitors. The virions were harvested from the supernatant by centrifugation through 200ul of 30% sucrose at 10,000 RPM at 4°C for 2hrs (Eppendorf Centrifuge 5424). The resulting pellet of virions was resuspended in 20ul of RIPA lysis buffer containing PR inhibitors.

### Co-assembly Experiments: Luciferase Assay

To test the infectivity of the virions produced from the HEK293 cells, TZM-bl cells were plated onto a 6-well tissue culture plate and incubated with 1ml of supernatant derived from HEK293 cells transfected with proviral DNA. These cells were then incubated at 37°C for 36-48 hrs. A britelite plus reporter gene assay system (Revvity) was then used to assess the number of cells that were infected by the virions.

### Co-assembly Experiments: Western Blotting

Cell extracts were centrifuged for 20 min at 15,000 RPM (Eppendorf Centrifuge 5424) before being diluted approximately 8X in RIPA lysis buffer. Virions were diluted approximately 2X in the same lysis buffer. Samples were then denatured in 10 ul of Lamelli sample buffer (Bio-Rad Laboratories) containing 5% BME and boiled at 95°C for 5min. The samples were loaded into Mini-PROTEAN TGX precast gels (Bio-Rad Laboratories) and their proteins were separated using SDS-PAGE. After completion of electrophoresis, the proteins were transferred onto a PVDF membrane (Millipore). This membrane was then washed with a blocking buffer (LI-COR Biosciences) and stained with anti-HIV-1 p24 monoclonal primary antibody (183-H12-5C, NIH HRP) overnight. An anti-mouse secondary antibody (LI-COR Biosciences) was then used to immuno probe the membrane for 45 min before it was scanned with the Odyssey Infrared Imaging System (LI-COR Biosciences) at 700 nm following manufacturer protocol.

### Coassembly Experiments: Monte Carlo Simulations

To calculate the infectivity of virions released for a range of NL4.3 to NL4.3:(mutant) ratios Monte Carlo simulation 10,000 cells were modeled. In each cell, the program assigned a total number of plasmids based on the distribution of number of plasmids from Figure 1. The following ratios of NL4.3/[NL4.3+NL4.3:(mutant)] were used (1, 0.9, 0.8, 0.7, 0.6, 0.5, 0.4 0.3, 0.2, 0.1, 0.08 0.07 0.06 0.05 0.04 0.03 0.02 0.01 0). In each cell, based on the assigned total number of plasmids to the cell, number of NL4.3 and NL4.3:(mutant) plasmids were assigned based on the above ratios. The program would run through all 10,000 cells for each one of the ratios presented above. Once the total number of plasmids and the number of the NL4.3 and NL4.3:(mutant) plasmids in the cell were assigned, the ratio of mutant and wild type proteins in the cytosol were put proportional to the number of NL4.3 and NL4.3:(mutant) plasmids respectively. It was assumed that each cell produces a set number of virions, for simplicity we assumed this number to be 1,000. The number of proteins in each virion was assumed to be “N” and for each virion, the program would randomly select proteins from the mix of wild type and mutant proteins in the cytosol. At the end, the program would have assembled 1,000 virions from each of the 10,000 cells modeled, which would make the total number of virions modeled to be 10 million. It is not known how the infectivity of each virion would vary based on the number of mutant versus wild type protein incorporated in the virion. We therefore used a minimum number of wild type proteins required for infectivity as “n”, which is unknown and needs to be fixed when fitting the model to the infectivity data. The Monte Carlo experiments as described above have only two free parameters, N: the total number of proteins of interest in the virion and n: the number of wild type proteins in each virion required for infectivity.

### Cell plating and transfections for counting the number of plasmids

HEK293 cells were plated onto 6-well tissue culture dishes with 2 mL of medium per well and incubated for 24 hours prior to transfection. At 50% confluency, the HEK 293 cells were co-transfected with NL4.3(iGFP)(D_25_N), NL4.3(imCherry)(D_25_N), and pNL4.3(D_25_N) at varying dilutions per well. Between the 3 plasmids, 4ng of plasmid (for each well of the 6 well dishes) were used for each transfection at dilutions between 1:2 and 1:256 for the imCherry and iGFP plasmids, with the remainder of the 4ng being pNL4.3(D_25_N) plasmid. The goal for using the different dilutions was to reach the stochastic limit for random distribution of the plasmids within the cells. Upon reaching said limit, a fraction of cells would only have one type of fluorophore coding plasmid and thereby only express a single-colored fluorophore. Transfection was carried out using 100 ul of Opti-MEM Reduced Serum Medium (Gibco) and 12 ul of Lipofectamine 2000 (Thermo Fisher Scientific) per well. Images were taken 24-48 hours post transfection using an Olympus CKX53 inverted microscope equipped with an Olympus EP50 camera, a brightfield illumination source, and a CoolLED pE-300 light source for red and green fluorescence excitation. All dilutions were imaged using brightfield, 488 nm, and 561 nm light sources to collect background images and individually excite iGFP and imCherry proteins.

### Measuring the probability of infection based on number of active Gag-Pol PR, RT and IN enzymes formed within HIV-1 virions

To determine how infectivity relates to the number of active enzymes in a virion, we utilized a stochastic sampling simulation. The simulation randomly selects two Gag-Pol molecules out of an assortment of NL4.3 and mutated Gag-Pols and identifies what combination has been selected. The selected pair of Gag-Pols is then removed from the selection pool. This is repeated until all pairs of Gag-Pols in the virion have been assessed. The whole process is done 200 times which yields enough data to visualize the distribution of active enzymes. The 3D plots of *Figure 4* present the distribution of active enzymes plotted against the percentage of NL4.3 Gag-Pol proteins.

The probability of being infectious plots (*f*_PR_(x), *f*_IN_(x), and *f*_RT_(x)) can be adjusted such that their x-axis is the percentage of NL4.3 Gag-Pol proteins. Doing so means that the probability of being infectious and the active enzyme results share a relationship with the percentage of NL4.3 Gag-Pol proteins. This relationship allows us to plot the probability of being infectious as a function of the mean number of active enzymes. The results of this are shown in the lower plots of *Figure 4*. According to this analysis HIV-1 virions reach a 50% chance of being infectious at 6 active PR enzymes and 99% at 11 active PR enzymes. RT reaches 50% at 39 active enzymes and 99% at 48 active enzymes.

## Author Contributions

GC performed infectivity experiments, analyzed data and performed modeling. HD harvested virions, performed infectivity experiments, cell culture, viral assembly, and western blot analysis, AD designed and ran drug infectivity experiments using robotics and optimized robotic readout and data analysis, WP harvested virions and performed infectivity experiments, BM measured virion infectivity, RG harvested virions and performed analysis utilizing monte carlo algorithm., NB performed and analyzed sequencing, BP performed cryotomography analysis, GY performed cell culture and viral assembly, AP performed sequencing, SS conceptualized the project, designed experiments and wrote the manuscript.

## Acknowledgments

Funding: This research was supported by NIH R56 AI150474-06A1 & NIH R01 AI186663-01 to (SS)

